# Genetic Basis for Lentil Adaptation to Summer Cropping in Northern Temperate Environments

**DOI:** 10.1101/2021.04.11.439349

**Authors:** Teketel A. Haile, Robert Stonehouse, James L. Weller, Kirstin E. Bett

## Abstract

The continued success of lentil (*Lens culinaris* Medik.) genetic improvement relies on the availability of broad genetic diversity and new alleles need to be identified and incorporated into the cultivated gene pool. Availability of robust and predictive markers greatly enhances the precise transfer of genomic regions from unadapted germplasm. Quantitative trait loci (QTLs) for key phenological traits in lentil were located using a recombinant inbreed line (RIL) population derived from a cross between an Ethiopian landrace (ILL 1704) and a northern temperate cultivar (CDC Robin). Field experiments were conducted at Sutherland research farm in Saskatoon and at Rosthern, Saskatchewan, Canada during 2018 and 2019. A linkage map was constructed using 21,634 SNPs located on seven linkage groups (LGs) which correspond to the seven haploid chromosomes of lentil. Eight QTL were identified for six phenological traits. Flowering related QTL were identified at two regions on LG6. *FLOWERING LOCUS T* (*FT*) genes were annotated within the flowering time QTL interval based on the lentil reference genome. Similarly, a major QTL for post-flowering developmental processes was located on LG5 with several senescence-associated genes annotated within the QTL interval. The flowering time QTL was validated in a different genetic background indicating the potential use of the identified markers for marker-assisted selection to precisely transfer genomic regions from exotic germplasm into elite crop cultivars without disrupting adaptation.

**Core Ideas:**

- Stable QTL were located for key phenological traits in lentil that lead to regional adaptation.
- *FT* genes are candidates for controlling flowering time in lentil grown in temperate environments.
- A major locus controlling post-flowering developmental processes was located on lentil LG5 with several senescence-associated genes annotated within the QTL interval.
- Markers identified in this study can be useful for marker-assisted selection to precisely transfer genomic regions from exotic germplasm into elite lentil cultivars without disrupting adaptation.

## INTRODUCTION

Lentil is a cool season food legume that plays a major role in alleviating food and nutritional insecurity. It also has significant agroecological value through its capacity for symbiotic fixation of atmospheric nitrogen and its contribution to management of pests and herbicide residue when grown in crop rotations. In 2019, lentil was produced in over 40 countries on over 4.8 million hectares globally (FAO, 2020). The top five lentil producing countries in 2019 were Canada, India, Australia, Turkey, and Nepal, with Canada being both the leading producer and exporter of lentil in the world (FAO, 2020). The increasing global demand for healthier, legume-based diets requires the continual development of widely adapted and higher yielding cultivars for diverse ecologies. Plant breeders often rely on elite breeding lines and cultivars for crossing because using unadapted germplasm may also introduce undesirable alleles, disrupting favorable allele combinations that have been fixed through several cycles of breeding. In many crops, elite breeding materials were derived from few founder lines and already have a narrow genetic base. Improvement of cultivars through conventional breeding techniques of selection-recombination-selection may stagnate if it is based on this narrow genetic diversity. The genetic diversity of modern lentil cultivars is relatively narrow compared to the diversity within the species and its wild relatives (Ahmad et al., 1996; Khazaei et al., 2016). Introducing new genetic variation from exotic sources is a crucial step to broaden the genetic base of modern lentil cultivars and to overcome challenges from new biotic and abiotic stresses.

Lentil originated in the Fertile Crescent of the Near East, which still harbors rich genetic diversity for both the cultivated primitive forms and wild relatives (Fratini et al., 2014; Lombardi et al., 2014). The spread of lentil from its center of origin to other parts of the world has been accompanied by the selection of traits which play a major role in adaptation to new agroecological zones (Materne & Siddique, 2009). Flowering, the transition from vegetative to reproductive stage, is one of the most important factors that determine the adaptation of plants to new environments. Other developmental processes such as stem elongation, apical dominance, lateral branching, resource allocation, maturity, and yield also accompany this transition (Weller & Ortega, 2015). Genes and environmental factors that affect the initial flowering transition can also influence post-flowering processes affecting fertility and pod development (Weller & Ortega, 2015). Lentil has a requirement of exposure to specific photoperiods and/or temperatures to flower, and flowering may be significantly delayed or prevented if these requirements are not met (Roberts et al., 1986; Summerfield et al., 1985a; Summerfield et al., 1985b; Wright et al., 2021). Lentils have been selected for adaptations to three major climatic regions of the world: Mediterranean, subtropical savannah (south Asia), and northern temperate (Materne & Siddique, 2009; Tullu et al., 2011). In Mediterranean-type environments, lentils are generally sown after the autumn rains and emerge into cool temperatures and short days, but vegetative crops experience progressively lengthening days and warming temperatures. In subtropical savannah environments, lentils are sown during the winter months and emerge into short days and cooler temperatures, but the temperature gradually rises during the reproductive phase. In northern temperate environments such as central Canada, lentil is usually seeded in the spring and plants grow and flower under long days and when temperatures are optimum. When germplasm from winter production regions is grown in northern temperate environments, flowering often occurs too early, before plants can accumulate the vegetative growth necessary to maximize their yield potential.

Identifying the genetic basis for flowering is crucial to develop genomic tools that will accelerate and enhance the precision of introducing new genetic variability from diverse germplasm. Several flowering-time genes that regulate the transition from vegetative to reproductive phase have been reported in crop and model legumes (Nelson et al., 2010; Weller et al., 2012; Weller & Ortega, 2015). This transition is triggered by environmental cues as well as endogenous signals such as developmental stage, hormones, and carbohydrate levels (Amasino & Michaels, 2010; Hecht et al., 2005). Light, through its effect on daylength and light-quality, and temperature, through vernalization and ambient temperature effects, are the main environmental regulators of flowering (Hecht et al., 2005). Endogenous pathways function independently of environmental signals and are related to the developmental state of the plant (Amasino & Michaels, 2010). In Arabidopsis (*Arabidopsis thaliana*), several genes influence the flowering process, with varied roles in photoperception, circadian clock function, photoperiod response, vernalization response, autonomous flowering, and integration of flowering pathway signals (Andrés & Coupland, 2012; Fowler et al., 1999; Imaizumi et al., 2003; Kardailsky et al., 1999; Kobayashi et al., 1999; Putterill et al., 1995; Searle et al., 2006; Simpson, 2004; Suárez-López et al., 2001). Many of these genes are shown to be conserved across legume crop species although several gene families have undergone differential gene loss or expansion (Hecht et al., 2005). For example, around 20 loci associated with flowering time have been reported in pea (*Pisum sativum*) and soybean (*Glycine max*) (Lin et al., 2021; Weller & Ortega, 2015). In chickpea (*Cicer arietinum*), four major genes involved in the control of flowering time have been identified (Gaur et al., 2015; Hegde, 2010; Kumar & van Rheenen, 2000). In lentil, Sarker et al. (1999) demonstrated the control of flowering time by the major locus *Sn*, which was subsequently identified as an *ELF3* ortholog (Weller et al., 2012). Beyond this, the genetic control of flowering time in lentil is not well understood. Several studies have reported flowering time QTL using intraspecific crosses (Fedoruk et al., 2013; Kahriman et al., 2015; Tullu et al., 2008) and interspecific crosses (Fratini et al., 2007; Polanco et al., 2019), but there is a notable lack of information on genetic differences that underlie major regional adaptations within the crop species. In this study, we used a RIL population developed from a cross between an exotic accession and adapted cultivar to identify regions of the lentil genome associated with key phenological traits. The availability of molecular markers for these loci is crucial for marker-assisted breeding and the effective use of broader lentil genetic resources in a breeding program.

## MATERIALS AND METHODS

### Plant materials and phenotyping

A population of 110 RILs was available from a cross between ILL 1704 and CDC Robin. CDC Robin is a lentil cultivar developed at the University of Saskatchewan (Vandenberg et al., 2002), and ILL 1704 is a landrace from Ethiopia. ILL 1704 was selected to represent the Mediterranean and northeast African lentil accessions. The RILs and parental lines were grown at Sutherland (lat 52°10’N, long 106°30’W) and Rosthern (lat 52°41’N, long 106°17’W), Saskatchewan in 2018 and 2019. All field plots were seeded between late April and mid-May with a target sowing density of 100 seeds per m^2^. The field trials were established in 1 m^2^ three-row microplots arranged in randomized complete block design with three replications in each site-year. Days to elongated tendrils (DTT), days to flowering (DTF), days to swollen pods (DTS), and days to maturity (DTM) were recorded in both years. DTT, DTF and DTS were recorded as the number of days from sowing to when 10% of plants in a plot had elongated tendrils, at least one open flower and pods with fully swollen seeds, respectively. Days to maturity was recorded as the date when 10% of plants displayed 50% pod maturity. Vegetative (VEG) and reproductive (REP) periods were recorded as the number of days from emergence to flowering and from flowering to maturity, respectively.

The phenotypic data were analyzed using analysis of variance with SAS mixed models, v9.4 (Littell et al., 2006). The data were first analyzed separately in each environment (site-year) and then combined across environments. Genotype was considered a fixed effect and replication was considered random for data analysis in each environment. For combined data analyses, replication nested in environment, environment, and genotype × environment interaction (GEI) were considered random effects. The Kenward-Roger degrees of freedom approximation method was used to compute the degrees of freedom for means (Kenward & Roger, 1997). Broad-sense heritability (H^2^) on a plot basis was estimated for all traits in each population using the equation 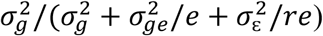, where 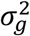 is the genetic variance, 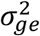 is the GEI variance, 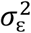 is the residual variance, *e* is the number of environments, and *r* is the number of replications per environment. Variance components were estimated in SAS using restricted maximum likelihood method described in Holland et al. (2003), with the effect of genotype, environment, GEI, and replication considered as random effect.

### Genotyping and QTL mapping

Genomic DNA was extracted from fresh leaves of 2 to 3-week-old seedlings for all lines using a Qiagen MiniPrep Kit (Qiagen). Genotyping of all lines was performed using a custom lentil exome capture assay as described in Ogutcen et al. (2018). Markers with a minor allele frequency higher than 5% and missing proportion less than 10% were used for linkage mapping. Markers showing significant segregation distortion (deviating from the expected 1:1 ratio for the RILs) based on a chi-square (χ^2^) test were removed from the analysis. We used a Perl pipeline, SimpleMap (Jighly et al., 2015), with a repulsion threshold of zero to check the number of recombination events for each pair of markers and to keep only one representative marker for pairs with no recombination. The reduced subset of representative markers was used for linkage mapping and redundant markers excluded at the first step were realigned to the map after linkage map construction (Jighly et al., 2015). Markers were clustered into linkage groups using the MSTMap software (Wu et al., 2008), with a *p*-value of 1E^-15^ and a maximum distance between markers of 15 cM. The relative order of markers within each linkage group was determined using the MapDisto v2.1.7.3 software (Lorieux, 2012), with a LOD threshold of 3 and a cut off recombination value of .3. Distances between markers were calculated using the Kosambi function (Kosambi, 1943). The order of markers in each linkage group was refined using AutoCheckInversions and AutoRipple commands in MapDisto. Linkage groups were corrected for double recombinants using the ‘color genotypes’ window in MapDisto. Linkage groups were assigned to each chromosome based on their marker locations on the lentil reference genome (CDC Redberry genome assembly v2.0) (Ramsay et al., 2019).

QTL analysis was performed using Windows QTL Cartographer v2.5 (Wang et al., 2012). Composite interval mapping (CIM) was implemented with a 2 cM walk speed. CIM was performed on the LS-means of each trait for individual environment and averaged (combined) across all environments. Cofactor selection was performed using forward and backward regression with a significance level of *p* = .1 with a 10 cM window size. QTL significance thresholds were determined based on 1000 permutations at a significance level of *p* = .05. Candidate genes within the identified QTL intervals were retrieved using the lentil reference genome (Ramsay et al., 2019).

### QTL validation

A total of 286 F_2_ lines were developed from a cross between CDC Redberry and ILL 1704. CDC Redberry is a lentil cultivar developed at the University of Saskatchewan (Vandenberg et al., 2006), and the genotype used for the reference assembly of the lentil genome. Both CDC Redberry and CDC Robin are adapted to northern temperate environments and share the same phenology characteristics; therefore, CDC Redberry is an acceptable alternative to CDC Robin for validating the identified QTL. In 2018, the 286 F_2_ lines and the two parents were grown at the Seed Farm of the Crop Development Center, University of Saskatchewan (lat 52°8’N, long 106°37’W). Each line was grown in 1.5-liter pots. Leaf tissues were collected from each 4 to 6-week-old seedling and plants were allowed to produce F_3_ seed. DNA was extracted using methods described above. In 2019, the resulting F_2_ derived F_3_ generations were grown as single 1 m rows at Sutherland adjacent to the LR-01 population. The field experiments were arranged in two blocks, each containing 51 plots. CDC Redberry, ILL 1704, CDC Kermit and CDC Maxim were grown as replicated checks. The field trials were established in 1 m^2^ three-row microplots arranged in an augmented randomized complete block design, where the four check cultivars were randomly assigned to rows of plots within a block and unreplicated entries were randomly arranged in the remaining rows (Federer, 1961). DTT, VEG, DTF, REP, DTM, and DTS were recorded as described above. The phenotypic data were analyzed using analysis of variance with SAS Mixed models, v9.4 (Littell et al., 2006). Genotypes (entries plus check cultivars) were considered as fixed effect and block was considered as a random effect. Trait values of entries were adjusted relative to the check cultivars replicated in each block using the LSMEANS procedure in SAS (Wolfinger et al., 1997).

A total of eight SNPs within the coding sequences of *FTb, FTa1, FTa2*, and *FTc* located in the flowering time QTL intervals were chosen for validation (Supplemental Table 1). Two allele specific forward primers and a reverse primer were designed for use in fluorescence based competitive allele-specific PCR assays (KASP; LGC Biosearch Technologies). DNA from the 286 lines was assayed using these primers and KASP reaction mix (LGC Biosearch Technologies) according to manufacturer’s instructions. PCR amplification was carried out in a CFX384 Real-Time PCR System (Bio-Rad) and end-product fluorescence readings were analysed using CFX Manager Software v3.1 (Bio-Rad).

## RESULTS

### Phenotypic data

Analysis of variance on combined data across environments revealed significant variation among the RILs for all measured traits (Table 1). There was no effect of the environment but the interaction of genotype with the environment was significant indicating variable performance of lentil genotypes across the four environments.

**Table 1.** *P* values from mixed model analysis of variance *F* test for six phonological traits.

| Effect | DTT | VEG | DTF | REP | DTM | DTS |
| --- | --- | --- | --- | --- | --- | --- |
| Genotype | *** | *** | *** | *** | *** | *** |
| Environment | NS† | NS | NS | NS | NS | NS |
| Replication (Environment) | * | * | * | NS | * | * |
| Genotype × Environment | *** | *** | *** | *** | *** | *** |
| CV% | 15.4 | 9.8 | 11.7 | 17.9 | 10.6 | 9.8 |

*Note:* DTT, days to elongated tendrils; VEG, vegetative period; DTF, days to flowering; REP, reproductive period; DTM, days to maturity; DTS, days to swollen pods.
\*Significant at the .05 probability level. \*\*\*Significant at the .001 probability level. †NS, nonsignificant.

Frequency distribution of traits for the RILs and means of the parental lines are shown in Figure 1. The distribution of traits followed an approximately normal distribution except for DTT and VEG at Sutherland in both years, VEG and DTF at Rosthern 2019, DTF at Sutherland 2018, REP and DTS at both locations in 2018, which were slightly skewed towards higher values, and DTM at Rosthern 2019, which was skewed towards lower values (Figure 1). As expected, ILL 1704 flowered and matured earlier than CDC Robin in all environments. ILL 1704 and CDC Robin differed in DTF by 4 to 9 days and in DTM by 7 to 15 days across environments. For all traits, transgressive segregants in both directions were observed in all environments (Figure 1).

**Figure 1.**
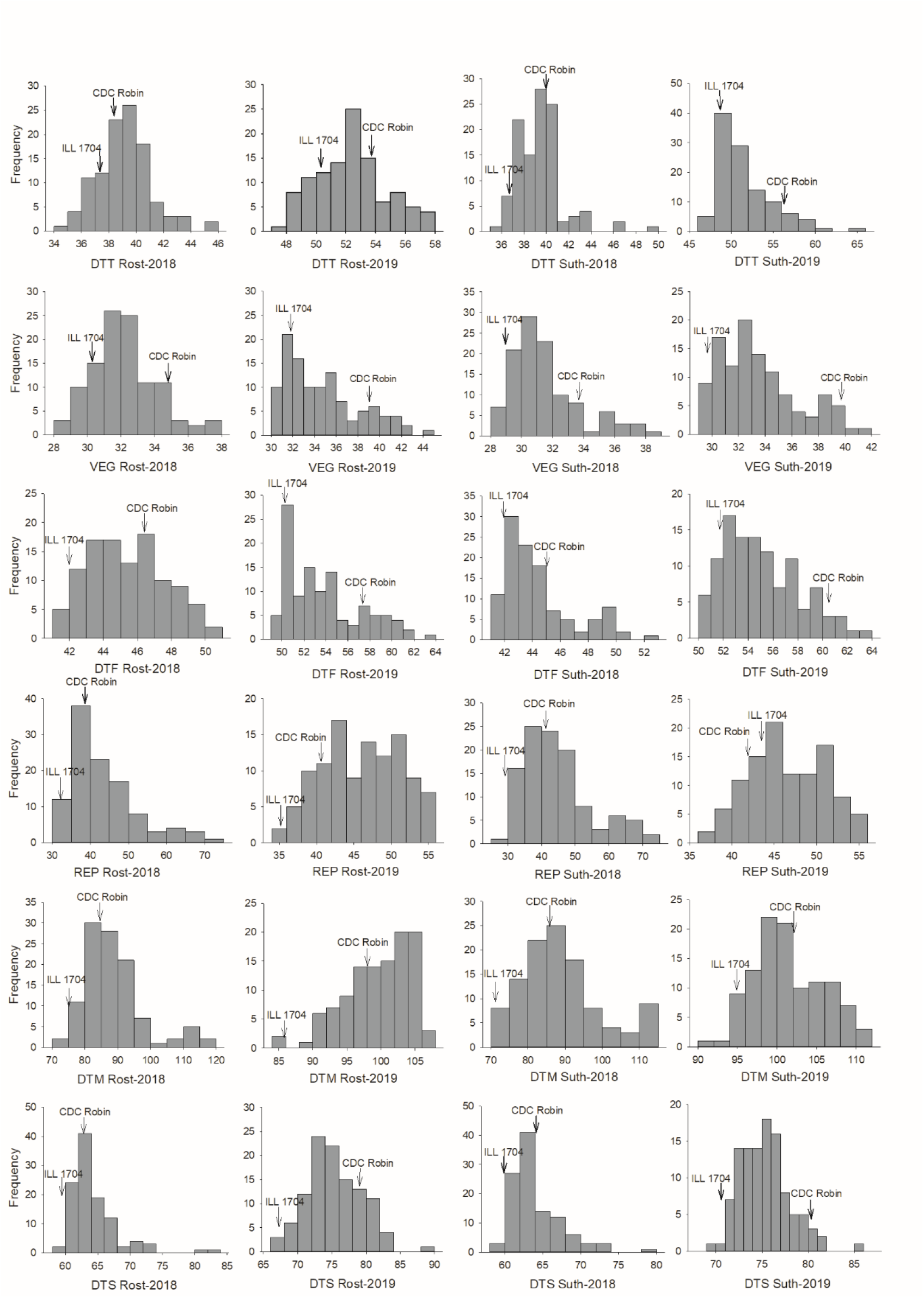
Frequency distribution of RILs and the two parents in the LR-01 population for days to elongated tendrils (DTT), vegetative period (VEG), days to flowering (DTF), reproductive period (REP), days to maturity (DTM), and days to swollen pods (DTS) measured at Rosthern (Rost) and Sutherland (Suth) in 2018 and 2019. The mean values of the parents are indicated with an arrow.

A considerable number of late transgressive segregants and very few early segregants were observed for DTM, REP, and DTS across environments. As these are essentially maturity rather than flowering traits it is likely that they are subject to more complex physiological and genetic control, exposing greater potential for the emergence of novel genotype combinations impairing maturity in some way. The two locations were more or less similar with respect to the performance of lines but there were slight differences between the years. In 2018, DTT ranged from 34 to 49 days across locations but it was delayed by up to two weeks in 2019. VEG was also delayed by up to a week in 2019 compared to 2018. Plants flowered earlier in 2018 compared to 2019 but flowering time was similar between Rosthern and Sutherland in both years. DTF ranged from 41 to 52 and 49 to 63 days in 2018 and 2019, respectively. Although the duration of flowering was earlier in 2018, maturity of plants took longer and as a result REP was narrower in 2019 at both locations.

Pearson’s correlation coefficients for all traits measured within each environment are shown in Table 2. There were significant positive correlations (.39 ≤ *r* ≤ .98, *P* < .001) among DTT, VEG, and DTF in all environments (Table 2). Similarly, DTM showed strong positive correlation with REP (.77 ≤ *r* ≤ .97, *P* < .001) and DTS (.42 ≤ *r* ≤ .83, *P* < .001) in all environments. DTM had no correlation with VEG but showed weak positive correlations with DTT and DTF across environments. DTS was also positively correlated to DTT (.36 ≤ *r* ≤ .75, *P* < .001) and DTF (.36 ≤ *r* ≤ .84, *P* < .001). On the other hand, REP showed either no correlation or weak to moderate negative correlations with DTT, DTF and VEG across environments. This may suggest distinct genetic control of flowering and maturity traits. Moderate to high estimates of heritability were obtained for DTT (.38), DTS (.49), DTF (.53), VEG (.58), REP (.61), and DTM (.62). The observed phenotypic variance in these traits has a genetic cause and the data can be successfully used to identify loci controlling phenological traits in lentil.

**Table 2.** Pearson’s correlation coefficients among six phenological traits for each environment.

| Trait | Environment | VEG | DTF | REP | DTM | DTS |
| --- | --- | --- | --- | --- | --- | --- |
| DTT | Rosthern 2018 | .39*** | .76*** | .04 | .23* | .36*** |
|  | Rosthern 2019 | .63*** | .68*** | -.11 | .35*** | .56*** |
|  | Sutherland 2018 | .54*** | .72*** | .16 | .31** | .56*** |
|  | Sutherland 2019 | .73*** | .82*** | -.29** | .28** | .75*** |
| VEG | Rosthern 2018 | — | .65*** | -.35*** | -.17 | -.03 |
|  | Rosthern 2019 | — | .98*** | -.54*** | .12 | .56*** |
|  | Sutherland 2018 | — | .90*** | -.10 | .07 | .30** |
|  | Sutherland 2019 | — | .92*** | -.47*** | .17 | .74*** |
| DTF | Rosthern 2018 | — | — | -.03 | .22* | .36*** |
|  | Rosthern 2019 | — | — | -.50*** | .17 | .61*** |
|  | Sutherland 2018 | — | — | -.01 | .21* | .46*** |
|  | Sutherland 2019 | — | — | -.43*** | .25* | .84*** |
| REP | Rosthern 2018 | — | — | — | .97*** | .76*** |
|  | Rosthern 2019 | — | — | — | .77*** | .20* |
|  | Sutherland 2018 | — | — | — | .97*** | .70*** |
|  | Sutherland 2019 | — | — | — | .77*** | -.17 |
| DTM | Rosthern 2018 | — | — | — | — | .83*** |
|  | Rosthern 2019 | — | — | — | — | .69*** |
|  | Sutherland 2018 | — | — | — | — | .79*** |
|  | Sutherland 2019 | — | — | — | — | .42*** |

\*Significant at the .05 probability level. \*\*Significant at the .01 probability level. \*\*\*Significant at the .001 probability level.

### Genetic map

The genetic map included 21,634 SNPs in seven linkage groups that spanned a total map distance of 1643.9 cM (https://knowpulse.usask.ca/Geneticmap/2695342; Supplemental Table 2). Linkage groups corresponded to the seven chromosomes of lentil (Supplemental Figure 2). For QTL analysis, SNPs with redundant positions were filtered out, leaving 2342 uniquely mapped SNPs (Supplemental Table 2). These were distributed evenly across chromosomes (from 228 in chromosome 7 to 421 in chromosome 3), with an average distance of .7 cM between adjacent SNPs.

### QTL analysis

QTL were only reported if they were identified in at least two of the individual environments and in the combined data, and across the six phenological traits only eight QTL met these criteria. Two QTL were detected each for VEG and DTF, and only one QTL each was detected for DTT, DTM, REP, and DTS (Table 3). The single QTL for REP, DTM, and DTS were localized within a common interval on LG5 (125.3-128.7 cM), whereas all remaining QTL were located within two distinct regions on LG6. QTL for DTT (*qDTT*.*6*), VEG (*qVEG*.*6-1*), and DTF (*qDTF*.*6-1*) were located near the top of LG6 (2.6-10.2 cM) and respectively explained 21-38%, 7-38%, and 19-43% of the phenotypic variance across environments (Table 3). The second region in the middle of LG6 (85.2-109.5 cM) also harbored QTL for VEG (*qVEG*.*6-2*) and DTF (*qDTF*.*6-2*) that accounted for 19-21% and 13-18% of the variance across environments, respectively. QTL for DTM (*qDTM*.*5*) and REP (*qREP*.*5*) explained 48 and 59% of the variance across environments, respectively. These QTL were also identified in all of the individual environments and explained 24 to 57% and 30 to 43% of the phenotypic variance, respectively. A single QTL for DTS (*qDTS*.*5*) was also localized with *qDTM*.*5* and *qREP*.*5* that accounted for 10 to 20% of the variance across environments.

**Table 3.** QTL identified for six phenological traits in the LR-01 population (ILL 1704 × CDC Robin) evaluated across four environments. QTL analyses were performed using data from each environment and averaged (combined) across all environments.

| QTL | Trait | Environment | LG | Position (cM) | Interval (cM) | LOD | R<sup>2</sup> (%) | ADD<sup>†</sup> |
| --- | --- | --- | --- | --- | --- | --- | --- | --- |
| *qDTT.6* | DTT | Combined | 6 | 6.8 | 6.3 – 8 | 14.2 | 33 | -1.19 |
|  |  | Rost-2019 | 6 | 7.8 | 7.4 – 8.3 | 8.2 | 22.5 | -1.13 |
|  |  | Suth-2018 | 6 | 6.8 | 6.3 – 8 | 10.2 | 21.1 | -.96 |
|  |  | Suth-2019 | 6 | 6.8 | 6.3 – 7.9 | 16.9 | 37.8 | -2.11 |
| *qVEG.6-1* | VEG | Combined | 6 | 4.8 | 4.3 – 5.3 | 13.9 | 27.6 | -1.33 |
|  |  | Rost-2018 | 6 | 4.8 | 3.3 – 5.7 | 4.2 | 7.3 | -.54 |
|  |  | Rost-2019 | 6 | 6.8 | 6.1 – 10.2 | 7.4 | 16.1 | -1.48 |
|  |  | Suth-2018 | 6 | 3.6 | 2.6 – 4.2 | 12.3 | 32.1 | -1.29 |
|  |  | Suth-2019 | 6 | 4.8 | 4.3 – 5.1 | 19 | 37.9 | -1.77 |
| *qVEG.6-2* | VEG | Combined | 6 | 91.3 | 90.7 – 93.2 | 10.2 | 18.7 | 1.07 |
|  |  | Rost-2019 | 6 | 85.4 | 85.2 – 85.9 | 8.3 | 18.5 | 1.56 |
|  |  | Suth-2018 | 6 | 102.6 | 101.6 – 103.7 | 8.6 | 18.8 | 1.04 |
|  |  | Suth-2019 | 6 | 89.1 | 88.5 – 90.2 | 12.5 | 21.1 | 1.34 |
| *qDTF.6-1* | DTF | Combined | 6 | 6.8 | 6.7 – 9.2 | 12 | 25.7 | -1.29 |
|  |  | Rost-2019 | 6 | 6.8 | 6 – 9.6 | 8.8 | 18.9 | -1.62 |
|  |  | Suth-2018 | 6 | 3.6 | 3 – 5 | 9.2 | 23.9 | -1.22 |
|  |  | Suth-2019 | 6 | 6.8 | 6.3 – 8.7 | 20.6 | 42.7 | -2.08 |
| *qDTF.6-2* | DTF | Combined | 6 | 104 | 102 – 104.5 | 6 | 10.9 | .94 |
|  |  | Rost-2018 | 6 | 104 | 103.1 – 104.7 | 5.7 | 13.2 | .96 |
|  |  | Rost-2019 | 6 | 90.7 | 88 – 91.9 | 8.6 | 18.1 | 1.55 |
|  |  | Suth-2018 | 6 | 92.3 | 91.5 – 94.2 | 7.1 | 17.1 | 1.06 |
|  |  | Suth-2019 | 6 | 106.6 | 105.8 – 109.5 | 8.4 | 13.6 | 1.13 |
| *qREP.5* | REP | Combined | 5 | 128.1 | 127.6 – 128.3 | 26.3 | 58.5 | -5.78 |
|  |  | Rost-2018 | 5 | 127.6 | 127.1 – 128.2 | 17.9 | 43.1 | -5.83 |
|  |  | Rost-2019 | 5 | 121.5 | 121 – 122.6 | 15.8 | 29.8 | -2.95 |
|  |  | Suth-2018 | 5 | 128.1 | 127.3 – 129.2 | 17.7 | 42.3 | -6.78 |
|  |  | Suth-2019 | 5 | 119.9 | 119.4 – 121.3 | 21.1 | 42.4 | -3.14 |
| *qDTM.5* | DTM | Combined | 5 | 128.1 | 127.2 – 128.7 | 19.7 | 48.4 | -5.02 |
|  |  | Rost-2018 | 5 | 128.1 | 127.4 – 128.5 | 15.3 | 39.1 | -5.72 |
|  |  | Rost-2019 | 5 | 127.6 | 127.2 – 128.2 | 16.7 | 40.4 | -3.00 |
|  |  | Suth-2018 | 5 | 127.1 | 126.1 – 128.3 | 11 | 24 | -5.64 |
|  |  | Suth-2019 | 5 | 128.1 | 127.6 – 128.4 | 24.3 | 55.6 | -3.53 |
| *qDTS.5* | DTS | Combined | 5 | 128.1 | 125.3 – 128.7 | 7.2 | 16.6 | -1.34 |
|  |  | Rost-2018 | 5 | 119.9 | 119.2 – 120.6 | 8.8 | 19.8 | -1.80 |
|  |  | Rost-2019 | 5 | 127.6 | 125.1 – 128.1 | 3.9 | 9.7 | -1.33 |
|  |  | Suth-2018 | 5 | 128.1 | 126.2 – 129.2 | 7 | 18 | -1.45 |

*Note:* DTT, days to elongated tendrils; VEG, vegetative period; DTF, days to flowering; REP, reproductive period; DTM, days to maturity; DTS, days to swollen pods; Rost, Rosthern; Suth, Sutherland; LG, linkage group; LOD, logarithm of the odds.
<sup>†</sup>ADD, additive effect attributed to this locus. Negative values represent alleles originating in ILL 1704; positive values represent alleles originating from CDC Robin.

## DISCUSSION

Control of flowering time is an important factor that determines plant adaptation to a new environment. The transition from vegetative to reproductive growth is a key developmental stage in the life of a plant, and the induction, expression, and maintenance of the flowering state are regulated by many external and endogenous factors (Hecht et al., 2005). In winter production regions, lentil is sown during winter months and emerges into short days and cooler temperatures. In Mediterranean-type environments, vegetative plants experience lengthening days and warmer temperatures slowly as the season progresses towards spring, whereas in subtropical savannah growing environments the temperature rises towards reproductive phase, but days are short during the entire crop life cycle. When lentils from these regions are sown in northern temperate environments during the spring, they experience long days and warm temperatures shortly after germination and their photoperiodic and critical temperature requirement for flowering are met quickly. Early flowering results in short stature plants that mature prematurely leading to reduced yield. Therefore, adaptations to delay flowering until plants accumulate enough biomass are necessary to avoid a significant yield penalty. Understanding the genetic control of flowering and post-flowering developmental processes is important to develop molecular tools that aid in germplasm evaluation as well as selection of adapted lines following hybridization with exotic germplasm.

We used a RIL population derived from a cross between a landrace from Ethiopia and a northern temperate cultivar to identify QTL associated with important phenological traits that lead to regional adaptations. Significant genetic variation was detected among the RILs for flowering and post-flowering developmental traits at the four environments (Table 1). Major QTL controlling DTT (*qDTT*.*6*), VEG (*qVEG*.*6-1*), and DTF (*qDTF*.*6-1*) were located near the top of LG6. *qDTT*.*6* and *qDTF*.*6-1* were co-localized while *qVEG*.*6-1* mapped 2 cM away which suggests a common genetic control for these correlated traits (Tables 2 and 3). This is expected because VEG is similar to DTF except that it is adjusted for days to emergence. Similarly, the development of tendrils preceded the transition from vegetative to reproductive stage and is correlated with flower initiation. As expected, the northern temperate cultivar contributed the positive alleles at these QTL that delayed the time to tendril development and flowering (Table 3).

The *qDTF*.*6-1* QTL interval spanned a 1.7 Mb region (from 1,050,443 to 2,724,479 bp) in the lentil reference genome v2.0 that harbors 25 annotated genes; including two genes homologous to the Arabidopsis florigen gene *FT (Lcu*.*2RBY*.*6g000730* and *Lcu*.*2RBY*.*6g000760)* which are directly under the QTL peak (Supplemental Table 3). Among the 585 SNPs within *qDTF*.*6-1* interval, ten are in the *Lcu*.*2RBY*.*6g000730* coding sequence and eleven within the *Lcu*.*2RBY*.*6g000760* coding sequence (Supplemental Table 4), and these may potentially serve as suitable markers for indicating the allelic state of *qDTF*.*6-1* locus.

In Arabidopsis, *FT* is expressed in phloem tissues of cotyledons and leaves under long days and encodes the hormone-like florigen protein that is transported through the phloem to the shoot apex where it promotes flowering (Abe et al., 2005; Wigge et al., 2005). The role of *FT* genes in the integration of environmental signals for flowering has been documented in a wide range of plant species including both model and crop legumes (Kong et al., 2010; Laurie et al., 2011; Yamashino et al., 2013). Natural variation for flowering time is associated with the corresponding syntenic genomic regions in several temperate legume species (Weller & Ortega, 2015). Papilionoid legumes have three distinct subclades of *FT* genes: *FTa, FTb*, and *FTc*, and in Medicago (*Medicago truncatula*) and pea, the *FTa* and *FTb* subclades are represented by two genes each, namely *FTa1/a2* and *FTb1/b2* (Hecht et al., 2011). In both species, *FTa1* plays an important role controlling flowering and integrating photoperiod, vernalization, and light quality signaling, while *FTb* genes are primarily implicated in the photoperiod response (Hecht et al., 2011; Laurie et al., 2011). The *qDTF*.*6-1* region is collinear with a region of Medicago chromosome 7 previously observed to also contain the two *FTb* genes *MtFTb1* and MtFTb2 (Hecht et al., 2011), and sequence relationships confirm that the lentil genes also belong to the *FTb* subclade. The fact that *qDTF*.*6-1* was identified under the long day growing environments in Saskatchewan is consistent with the possibility that one or both of these genes may also have a role in promoting flowering under long day conditions and show attenuated function in genotypes adapted to the Canadian summer cropping system.

The second QTL for DTF (*qDTF*.*6-2*) on LG6 was characterized by alleles associated with reduced flowering time in the northern temperate cultivar (Table 3). The *qDTF*.*6-2* interval spanned 6.4 Mb in the lentil reference genome and harbors 96 annotated genes, including orthologs of the *FTa1, FTa2*, and *FTc* genes described previously in Medicago and pea (Hecht et al. 2011; Supplemental Table 3). Similarly, Ortega et al. (2019) reported the presence of a cluster of *FTa1, FTa2*, and *FTc* under the peak of a major flowering time QTL on chickpea chromosome 3. In Medicago and pea, significant roles have been reported for *FTa1* based on analyses of induced mutants (Hecht et al. 2011, Laurie et al. 2011). Mutants of both species are late-flowering under both long and short-days and in Medicago, transgenic overexpressing lines flower early with reduced photoperiod and vernalization requirements. The expression of *MtFTa1* corresponds with conditions that promote flowering in Medicago (Laurie et al., 2011). *MtFTa1* is up-regulated after a period of extended cold followed by exposures to warm long day photoperiods indicating that it integrates vernalisation and photoperiod signals.

In contrast, *MtFTa2* is expressed at higher levels in short rather than long days and is up-regulated during exposure to cold (Laurie et al., 2011). In pea, *FTa1* gene corresponds to the *GIGAS* locus, which encodes a mobile floral signal that is essential for flowering under long days and promotes flowering under short days but is not required for the photoperiodic response (Hecht et al., 2011). There are inconsistent reports about the role of vernalization on flowering in lentil. Summerfield et al. (1985b) evaluated the effects of temperature and photoperiod on six lentil genotypes and reported that all flowered sooner when grown from vernalized seeds, but genotypes differed in their relative sensitivities. On the other hand, Roberts et al. (1988) used two lentil genotypes, one of which was also used in Summerfield et al. (1985b) and reported no evidence of vernalization response in either cultivar. The *FTa* homologue in lentil may or may not be involved with vernalization response, however, at both Rosthern and Sutherland, night temperatures were around or below freezing for several days after sowing and the vernalization requirements of seeds, if any, would have been met. Unlike *FTa* and *FTb* which are expressed in leaves, *FTc* is only expressed in the shoot apex and plays a role in integration of signals from leaf-expressed *FT* genes (Hecht et al., 2011; Laurie et al., 2011). In chickpea, elevated expression of *FTa1* and *FTc* were consistently noticed in early flowering parents before floral induction under both short and long days but *FTa2* transcript was not detected in some of the early parents suggesting that *FTa1* and *FTc* are the likely candidates underlying their flowering time QTL (Ortega et al., 2019). Moreover, transgenic overexpression of Medicago and pea *FTa2* gene in Arabidopsis *ft-1* mutants showed weak activity to induce flowering compared to *FTa1* and *FTc* (Hecht et al., 2011; Laurie et al., 2011). This evidence suggests that both *FTa1* and *FTc* are the likely candidate genes at *qDTF*.*6-2* but it is also possible that the lentil *FT* genes have different expression or function. There are four, two, and six SNPs annotated within *FTa1, FTa2*, and *FTc* coding sequences in the lentil reference genome, respectively (Supplemental Table 4). These markers can be used to identify the allelic state of the *FTa* and *FTc* genes in lentil.

Previous reports of QTL associated with flowering time have linked one gene, *Sn*, to a seed coat pattern gene (*Scp*) (Fratini et al., 2007; Kahriman et al., 2015; Sarker et al., 1999). *Sn* was later revealed to be an *ELF3* ortholog (Weller et al., 2012) and it is found on chromosome 3 of the lentil reference assembly (Ramsay et al., 2019), so is not one of the genes responsible for variation in DTF in the LR-01 population. Results of mapping in a RIL population derived from a cross between a *L. culinaris* cultivar and a *L. orientalis* genotype also indicated linkage between a flowering QTL and seed coat pattern (Fratini et al., 2007), which led the authors to speculate that *Sn* was responsible. Fedoruk et al. (2013), however, mapped a seed coat pattern locus on chromosome 6. There were no DTF QTL near this locus in their temperate × temperate population; likely because those genes carried the same allele in both parents and, therefore, were not segregating. Using an interspecific cross between a *L. culinaris* cultivar and an *L. odemensis* accession grown in a short-day environment, Polanco et al. (2019) reported a major QTL for flowering time on chromosome 6 that explained 56 % of the observed variability. They also mapped a seed coat pattern locus nearby. Recently, our group identified a marker associated with seed coat pattern in a lentil diversity panel which maps to 12,192,948 bp region of chromosome 6 in the lentil reference genome (Wright & Bett, unpublished data, 2021). This suggests that the major flowering time QTL reported in Polanco et al. (2019) and *qDTF*.*6-1* identified in our study are likely the same and *FTb* may have a role in controlling flowering in both the long day and short day growing environments.

Given that there are at least two loci involved in seed coat pattern and the one on chromosome 6 is not linked to *Sn*, caution should be taken in interpreting the genetics of flowering time based on seed coat pattern. In pea, there are several known genes related to seed coat pattern (Blixt, 1974). One, the marbling locus *Marmoreus* (*M)*, is located near the *High response* (*Hr*) flowering time gene (Murfet, 1973). The identity of pea *Hr* and lentil *Sn* as orthologs of *ELF3* (Weller et al., 2012) suggests that the lentil *Scp* locus linked to *Sn* (Sarker et al., 1999) likely corresponds to the pea *M*, or possibly *F*, locus. Another pea locus, *Fs*, maps to a genomic region that is syntenic with the top of lentil chromosome 6 (Lamprecht, 1942a; Lamprecht, 1942b; Lamprecht, 1948; Sindhu et al., 2014), and could be the ortholog of the second lentil pattern gene. A single major QTL on LG5 was shown to affect REP, DTM, and DTS in LR-01 (Table 3).

This region is clearly important for controlling post-flowering developmental processes in lentil. The northern temperate cultivar contributed the alleles at this locus that delayed REP, DTM, and DTS (Table 3). There are a total of 69 SNPs within the QTL interval which spanned a large physical distance of 62.2 Mb (356,415,824 to 418,589,260 bp) in the lentil reference genome. Three other SNPs from the unanchored *unitig2307* were also located within the QTL interval. There are 561 annotated genes within the confidence interval of the QTL, which included, among others, embryonic abundant proteins, dehydration-responsive element-binding protein, and senescence-associated proteins (Supplemental Table 3). There are several potentially phenology-or senescence-associated genes in this broad interval and further research is required to narrow down the list.

Matching crop phenology to the growing environment is essential. In general, pulse breeding programs characterize breeding material by scoring DTF to identify lines that flower within the optimum window because measurement of DTM can be unreliable due to the occurrence of disease and adverse environmental conditions late into the growing season (Tullu et al., 2008). Availability of robust and predictive markers enhances the precision of identifying locally adapted germplasm.

In spring-summer cropping systems of the temperate regions, where temperature and photoperiodic requirements for floral induction are met early in the season, lentil breeding programs should consider selecting for the late alleles at *qDTF*.*6-1, qDTF*.*6-2*, and *qDTM*.*5*. This ensures that plants flower within the optimum window after accumulation of sufficient biomass leading to greater yield. In winter cropping systems such as Mediterranean and subtropical savannah production regions, photoperiodic and temperature requirements for flowering are met slowly as the season progresses but lentil production is limited by high temperature and terminal drought in spring and early summer (Nelson et al., 2010). Early flowering is a desirable trait in these environments and the early allele at these QTL may be worth pursuing. In our study, DTF showed weak positive correlations with DTM across environments (Table 2). Early flowering leads to early onset of reproductive growth but genotypes that flower early did not necessarily mature early and genotypes that flower late can have a short reproductive period and mature together with the ones that flowered within the optimum window. Therefore, combined selection for DTF and DTM at the QTL identified in our study would enable accurate prediction of genotypes that are adapted to a particular environment.

### Validation of identified QTL

We used an F_3_ generation from a cross between a temperate cultivar CDC Redberry and ILL 1704 for validating the flowering time QTL identified in this study. The distribution of phenological traits for the F_3_ and the parental lines are shown in Figure 2. Transgressive segregants in both directions were observed for all traits. The distribution of DTT, VEG, DTF, and DTS were slightly skewed toward higher values while DTM and DTS were skewed towards lower values (Figure 2). ILL 1704 flowered 10 days earlier and matured 11 days earlier than CDC Redberry.

**Figure 2.**
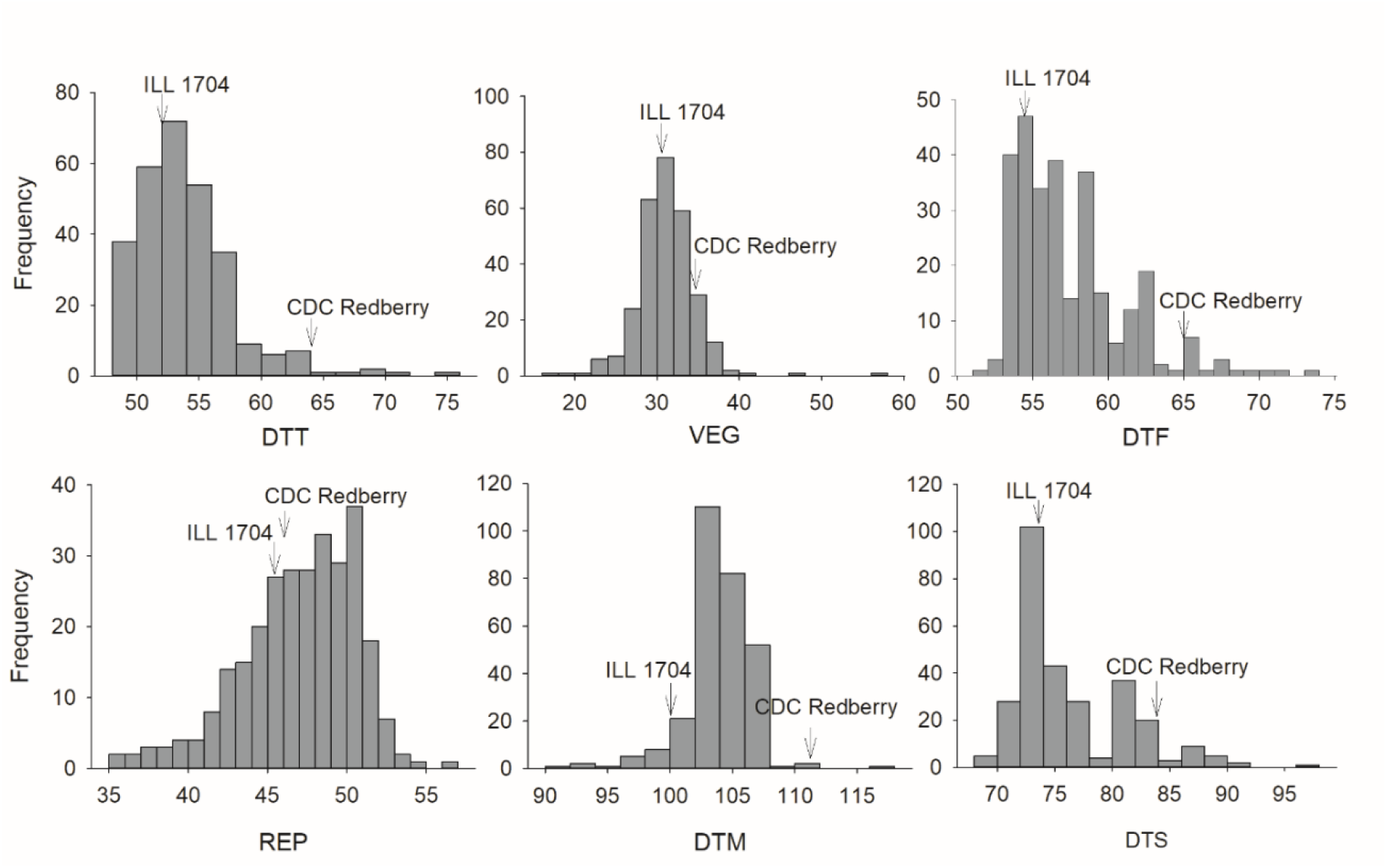
Frequency distribution of CDC Redberry × ILL 1704 F_3_ generation and the two parents for days to elongated tendrils (DTT), vegetative period (VEG), days to flowering (DTF), reproductive period (REP), days to maturity (DTM), and days to swollen pods (DTS). The mean values of the parents are indicated with an arrow.

Two SNPs within *FTb* coding sequence (*qDTF*.*6-1* interval) and six SNPs within *FTa1, FTa2*, and *FTc* coding sequences (*qDTF*.*6-2* interval) were converted to KASP primer and 286 lines were screened for DTF (Table 4; Supplemental Table 1). The SNPs within *FTa1, FTa2*, and *FTc* have similar haplotypes which suggests that the three genes originated from recent duplication events (Supplemental Table 1). Student’s *t-*test was performed to determine statistical significance of the difference in DTF between sets of individuals carrying the CDC Redberry and ILL 1704 alleles. DTF was significantly different (*t* = 2.3, *p* = .023) and (*t* = -2.6, *p* = .010) between the two alleles for both groups of markers. At *qDTF*.*6-1*, the average number of days to flowering were 58.1 (±3.6) and 56.6 (±4) between lines carrying the CDC Redberry and ILL 1704 alleles resulting in mean difference of 1.5 days. Similarly, mean number of days to flowering were 56.1 (±2.8) and 57.9 (±5.5) between lines carrying the CDC Redberry and ILL 1704 alleles at *qDTF*.*6-2* resulting in mean difference of 1.8 days. This suggests that the identified QTL can be useful in different genetic backgrounds involving crosses between adapted and exotic germplasm.

**Table 4.** Markers selected for validation of the flowering time QTL using an F_3_ generation developed from CDC Redberry × ILL 1704 cross.

| SNP | QTL | Location | Allele† |  | *t*-value | Pr > \|*t*\| |
| --- | --- | --- | --- | --- | --- | --- |
|  |  |  | CDC Redberry | ILL 1704 |  |  |
| *Lcu.2RBY.Chr6:2628041*<br>*Lcu.2RBY.Chr6:2528796* | *qDTF.6-1* | *FTb* | 58.1 (3.6) | 56.6 (4) | 2.3 | .023 |
| *Lcu.2RBY.Chr6:309231659*<br>*Lcu.2RBY.Chr6:309230402* | *qDTF.6-2* | *FTa1* | 56.1 (2.8) | 57.9 (5.5) | -2.6 | .010 |
| *Lcu.2RBY.Chr6:309276496*<br>*Lcu.2RBY.Chr6:309247768* | *qDTF.6-2* | *FTa2* | 56.1 (2.8) | 57.9 (5.5) | -2.6 | .010 |
| *Lcu.2RBY.Chr6:310035950*<br>*Lcu.2RBY.Chr6:310035979* | *qDTF.6-2* | *FTc* | 56.1 (2.8) | 57.9 (5.5) | -2.6 | .010 |

†Mean and standard deviation of average days to flowering between lines carrying CDC Redberry and ILL 1704 alleles.

In summary, QTL for key phenological traits in lentil were located using a RIL population derived from a cross between an exotic landrace and adapted cultivar. Two regions on LG6 harbored flowering-related QTL. Similarly, a stable region on LG5 was consistently associated with post-flowering developmental processes. The peak markers linked to the QTL fell either within the gene coding sequences or close to known genes controlling flowering time and senescence and can be used for marker-assisted breeding to transfer desirable alleles from exotic germplasm into elite lentil cultivars.

## ACKNOWLEDGEMENTS

This research was conducted as part of the “Application of Genomics to Innovation in the Lentil Economy (AGILE)”, a project funded by Genome Canada and managed by Genome Prairie. We are grateful for the matching financial support from the Saskatchewan Pulse Growers, Western Grains Research Foundation, the Government of Saskatchewan, and the University of Saskatchewan. We acknowledge the technical assistance of the bioinformatics, field and molecular lab staff of the Pulse Crop Breeding and Genetics group at the University of Saskatchewan.

## SUPPLEMENTAL MATERIAL

Supplemental Table 1 is a list of SNPs selected from the coding sequences of *FTb, FTa1, FTa2* and *FTc* that are annotated within the flowering time QTL intervals and used for validation of these QTL in an independent F_3_ population (CDC Redberry × ILL 1704). Supplemental Table 2 is a summary of the SNP based linkage map for LR-01 population (ILL 1704 × CDC Robin) used for QTL analyses. Supplemental Table 3 lists candidate genes annotated within the QTL intervals retrieved from the lentil reference genome (CDC Redberry genome assembly v2.0). Supplemental Table 4 lists all SNPs located within the coding sequences of the respective *FT* genes annotated within the flowering time QTL intervals in the lentil reference genome v2.0. Supplemental Figure 1 is the linkage map for the LR-01 population.

## Abbreviations

CIM: composite interval mapping
DTF: days to flowering
DTM: days to maturity
DTS: days to swollen pods
DTT: days to elongated tendrils
GEI: genotype × environment interaction
LG: linkage group
LOD: logarithm of the odds
QTL: quantitative trait loci
REP: reproductive period
RIL: recombinant inbreed line
VEG: vegetative period

