## Supplemental Table 2 for "Genetic Basis for Lentil Adaptation to Summer Cropping in Northern Temperate Environments"

**Supplemental Table 2.** Summary of the SNP based linkage map for LR-01 population (ILL 1704 × CDC Robin) used for QTL analyses.

| Linkage groups | Length (cM) | Number of SNPs | SNP density† | Maximum gap (cM) | Number of uniquely mapped SNPs |
|---|---|---|---|---|---|
| LG1 | 200.6 | 2806 | 14.0 | 4.7 | 269 |
| LG2 | 250.1 | 5120 | 20.5 | 3.5 | 393 |
| LG3 | 299.5 | 4540 | 15.2 | 6.5 | 421 |
| LG4 | 271.2 | 3563 | 13.1 | 4.8 | 377 |
| LG5 | 197.4 | 2004 | 10.2 | 3.3 | 279 |
| LG6 | 260.5 | 1891 | 7.3 | 2.9 | 375 |
| LG7 | 164.6 | 1710 | 10.4 | 2.9 | 228 |
| Whole genome | 1643.9 | 21634 | 13.2 | 6.5 | 2342 |

†Average number of SNPs per cM
