## Supplemental Figure 1 for "Genetic Basis for Lentil Adaptation to Summer Cropping in Northern Temperate Environments"

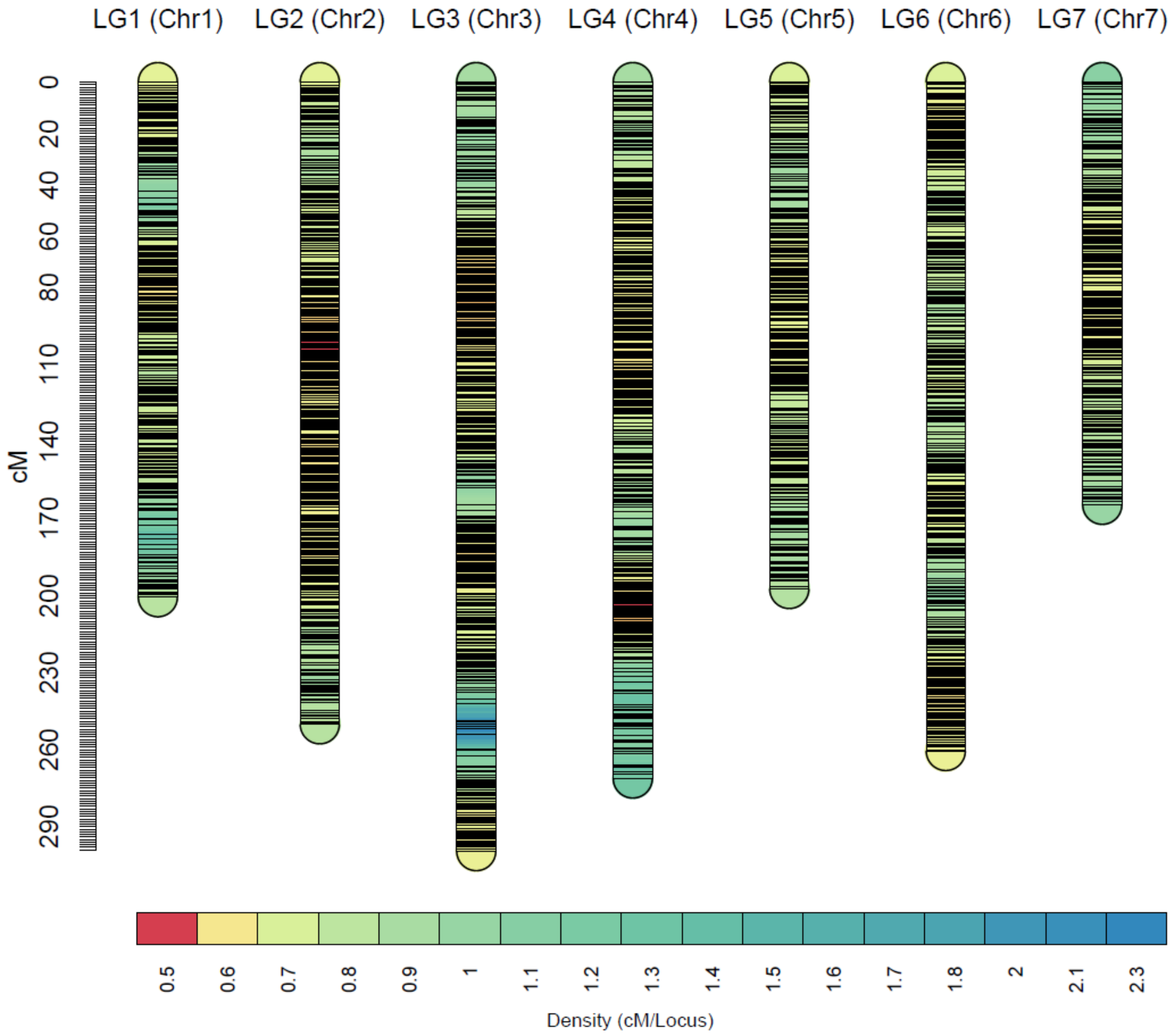


**Supplemental Figure 1**. High-density linkage map of LR-01 population (ILL 1704 × CDC Robin). The color bar shows the density of markers and horizontal lines in each linkage group indicate the position of markers.
